# The effect of histone deacetylase inhibitor Trichostatin A on IL-17A-induced transformation of MRC5 into myofibroblasts and its mechanism

**DOI:** 10.1101/2021.03.31.437856

**Authors:** Qian Zhang, Zexin Guo, Jing Zhang, Wei Zhou, Yueqing Wu, Lixia Dong, Jing Feng, Shuo Li, Jie Cao

## Abstract

**Objective:** To elucidate potential IL-17A- and TSA-mediated regulation of fibroblasts transformation.

**Methods:** MTT assay, HDAC1 activity assay, cell immunofluorescence and Western blot were employed to detect the expression of related indicators.

**Results:** MRC5 cells expressed only a small amount of Vimentin. IL-17A treatment upregulated MRC5 cell proliferation, in a concentration-dependent manner. TSA treatment, however, suppressed MRC5 cell proliferation. IL-17A treatment also upregulated HDAC1 activity in MRC5 cells, in a concentration-dependent manner. Using immunofluorescence, we demonstrated that IL-17A-treated MRC5 cells had markedly elevated Vimentin, Collagen-I and a-SMA levels, compared to controls. However, a combined treatment of IL-17A and TSA resulted in markedly reduced levels of the Vimentin, Collagen-I and a-SMA, compared to IL-17A alone, yet the amount was higher than controls. Using western blot analysis, we also revealed that the IL-17A-treated MRC5 cells had markedly elevated levels of Vimentin, a-SMA, HDAC1, p-Smad2, and p-Smad3, and markedly reduced level of Smad7, compared to controls. In TSA intervention group, the expression effect of the above protein was opposite. Moreover, no discernible difference was observed in the levels of Smad2 and Smad3 among the treated and un-treated groups.

**Conclusion:** IL-17A stimulates proliferation of MRC5 cells and increases HDAC1 activity and protein expression. It also transforms MRC5 cells into myofibroblasts via activation of the TGF -β1/ Smads signaling network. TSA, on the other hand, strongly suppresses TGF -β1/ Smads pathway-mediated fibrosis by ceasing HDAC1 activity and protein expression.

## Introduction

Incidences of idiopathic pulmonary fibrosis (IPF) are the most prevalent among interstitial lung diseases. Its etiology and pathogenesis are currently unknown. IPF has the characteristics of chronic, progressive and irreversible disease progression, with main pathological features being fibroblasts proliferation and aggregation, collagen fibers deposition in the lung interstitium, and activation and infiltration of lymphocytes, macrophages and monocytes, resulting in the devastating destruction of the alveolar structure. At present, clinical therapeutics for IPF include antioxidants, glucocorticoids, immunosuppressants, new anti pulmonary fibrosis drugs (such as: pirfenidone, nidanib), etc. Unfortunately, the above-mentioned drugs, whether used alone or in combination, does not significantly alter patient prognosis. In most cases, pulmonary fibrosis continues and the patient dies. Therefore, it is both urgent and necessary to examine the molecular mechanism of pulmonary fibrosis occurrence and development and simultaneously search for novel targets for IPF therapy. Emerging reports suggest a role for histone deacetylase (HDAC) in tumor development and organ fibrosis. Trichostatin A (TSA) is a commonly used HDAC inhibitor (HDACi). However, its mechanism of action in IPF remains unclear. It is crucial to determine whether TSA can alleviate pulmonary fibrosis via suppression of TGF - β1/Smads signaling network and also compare its affects on IPF with traditional anti-fibrosis drugs. The goal of this study is to explore a novel therapeutic target for IPF. We have, therefore, examined the effect of TSA on IL-17A-induced occurrence and development of pulmonary fibrosis in the human lung fibroblasts 5 (MRC5) cells.

## 1. Materials and methods

### 1.1 Reagents

MRC5 cells were donated to the school of life sciences, Tianjin NanKai University, Tianjin, China. IL-17A was purchased from Sino biological. TSA purchased from MedChemexpress. HDAC1 kit was purchased from Shanghai Jiemei Gene Medicine Technology. The MTT kit was purchased from Beijing Solebo Technology. HDAC1, Vimentin, Anti-a-SMA, anti-HDAC1, anti-p-Smad2, anti-p-smad3, anti-Smad2 and anti-Smad3 were acquired from Cell Signaling Technology of USA. Anti-collagen-I and anti-Smad7 were purchased from Abcam. Olympus cell sensentry was purchased from Olympus in Japan.

### 1.2 Methods

#### 1.2.1 Cell culture

MRC5 cells were grown in DMEM medium with 10% fetal bovine serum and in a humidified environment containing 5% CO2 at 37 °C. The cells were passaged at a cell density of ~90%. During passaging, the cells were trypsin-digested with 0.25% EDTA and the digestion was terminated in complete medium. Only cells that fell within the logarithmic growth phase were selected for experimentation.

#### 1.2.2 MTT assay for the detection of cell proliferation in IL-17A- and TSA- treated MRC5 cells

3×10^3^ cells per well were seeded in 96 well plates. The experimental setup included IL-17A-treated, TSA-treated, control (with cells and culture medium), and blank (with culture medium only). Varying concentrations of IL-17A were tested, namely, 25ng/ml, 50ng/ml, 100ng/ml, 200ng/ml, 300ng/ml, 400ng/ml and 500ng/ml. Similarly, varying concentrations of TSA were also examined, namely, 100nmol/L, 200nmol/L, 300nmol/L, 400nmol/L, 500nmol/L, 600nmol/L, 700nmol/L, 800nmol/L, 900nmol/L and 1000nmol/L. After intervention for 24, 48 and 72 hours, MTT reagent was introduced to the cells followed by further incubation for 4 hours. Absorbance was recorded at 490nm using an enzyme reader. Subsequently, cell proliferation was assessed as follows: Relative cell viability (%) = (control group of drug treatment group OD) / (control group of zero adjustment group OD)×100%.

#### 1.2.3 The IL-17A-mediated regulation of HDAC1 activity in MRC5 cells by the HDAC1 Kit

MRC5 cells were assigned to two groups. In one group, the nucleus was extracted by the nuclear extraction kit. In another, the HDAC activity was determined with the HDAC activity test kit, following operator’s guidelines. Finally, fluorescence intensity was detected by fluorescence microscope and the HDAC activity was calculated as follows: HDAC1 specific activity = sample total activity - Sample nonspecific activity.

#### 1.2.4 Cell immunofluorescence (IF) staining

1×10^5^/L MRC5 cells were seeded on glass slides. Upon adherence to the slide, they were PBS-washed three times and fixed with 1ml 40 g/L paraformaldehyde at room temperature (RT) for 20 min, sealed with goat serum blocking solution for 1 h, PBS-washed again three times, exposed to primary antibody (1:100) at 4 °C overnight (O/N), PBS-washed three times, exposed to secondary antibody (1:100) at RT for 1 h, PBS-washed three times, exposed to fluorescence reagent at RT for 1 h, PBS-washed three times, lastly, the anti fluorescence quenchant solution with DIPA was used to seal the film and images were taken under a laser confocal microscope.

#### 12.5 Evaluation of relevant proteins by Western Blot (WB)

Amount? MRC5 cells were seeded and cultured for 48 h. Cell lysates containing phosphatase inhibitor was centrifuged at 12000 rpm for 10 minutes and the supernatant was retrieved for protein quantification by BCA method. 20 ug of total protein was subsequently used for electrophoresis (SDS-PAGE), transferred into PDVF membrane, blocked with 5% skim milk at RT for 90 min, exposed to primary antibody O/N at 4 °C, rinsed 5 times with TBST, exposed to secondary antibody at RT for 60 min, TBST-rinsed 5 times, and exposed to hypersensitive ECL luminescent solution before obtaining images of proteins using a gel imaging system. GAPDH was used as internal reference. Image J software was used to calculate relative expression of target proteins.

### 1.3 Statistical methods

GraphPad software was employed for all statistical analyses. Data are presented as mean ± standard deviation. Pairwise comparison was done with t-test. P < 0.05 was the statistically significant threshold.

## Result

### 1. Identification of MRC5 cells

Our observations revealed that the MRC5 cells grew in a spindle form. The Vinmentin protein, which exhibited a green fluorescence, was localized in the cytoplasm. Alternately, the α - SMA protein, which also fluoresced green, was scarcely present in these cells. The nucleus was stained with fluorescent blue DAPI staining. Based on these observations, we confirmed, using biological and phenotypic characteristics of the cells, that the MRC5 cell line was functionally and phenotypically fibroblasts. (Figure 1)

**Fig. 1.**
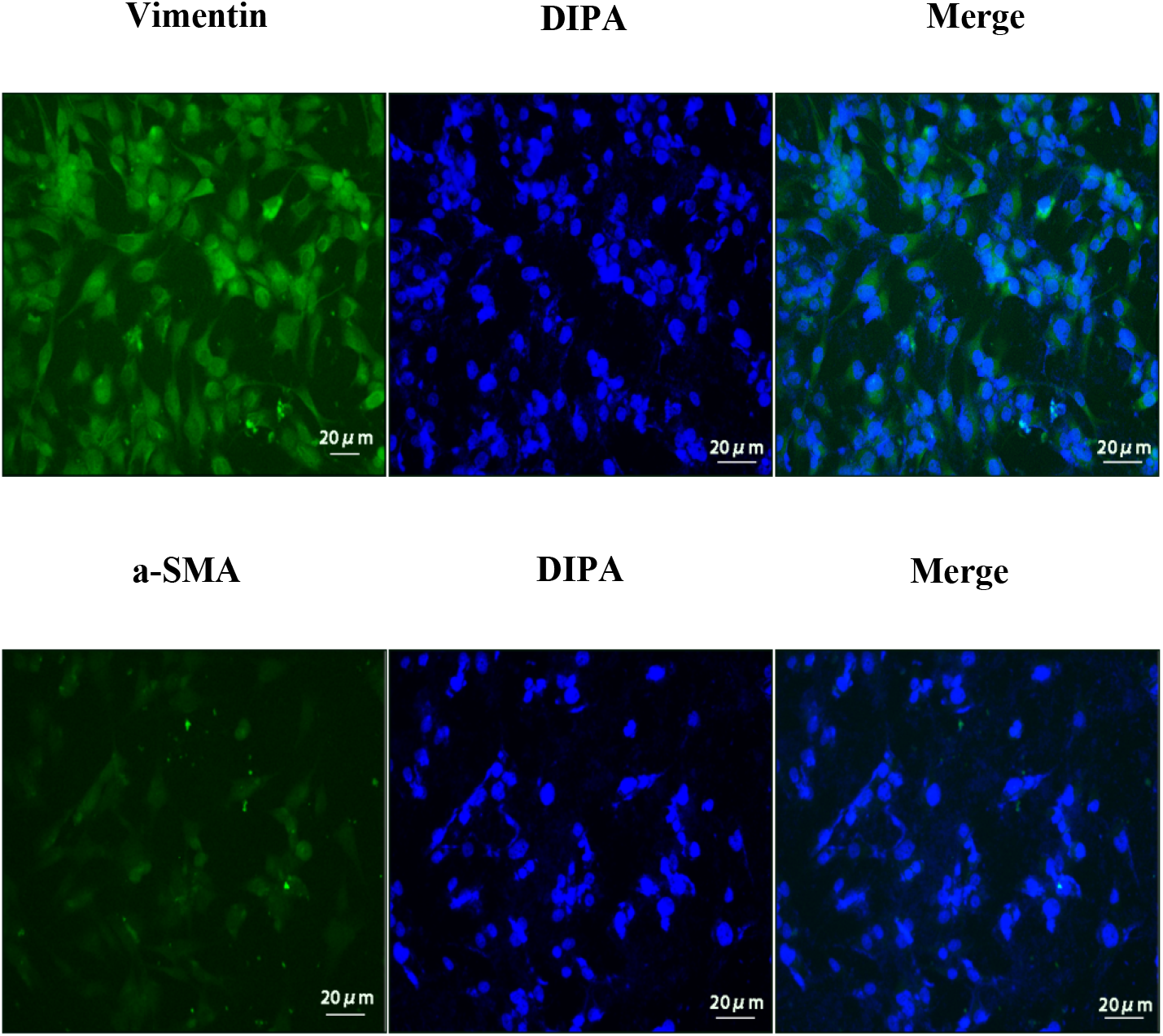
Immunofluorescence staining of MRC5 cells

### 2. Effects of differing concentrations of IL-17A on MRC5 cell proliferation

MTT assay revealed that IL-17A induced MRC5 cell proliferation, in a concentration dependent manner (p<0.05). In particular, 50 ng/ml IL-17A-treated cells had more proliferation than non-treated MRC5 cells (*aP*< 0.05). 300 ng/ml had higher proliferation than the 50 ng/ml IL-17A-treated MRC5 cells (*bP*< 0.05). Lastly, the 500 ng/ml IL-17A-treated cells had higher proliferation activity than the 300 ng/ml-treated MRC5 cells (*cP*< 0.05). Based on the results of our evaluation and the reference concentration in literature, 300 ng/ml IL-17A was selected for use in subsequent experimentations (Figure 2).

**Fig. 2.**
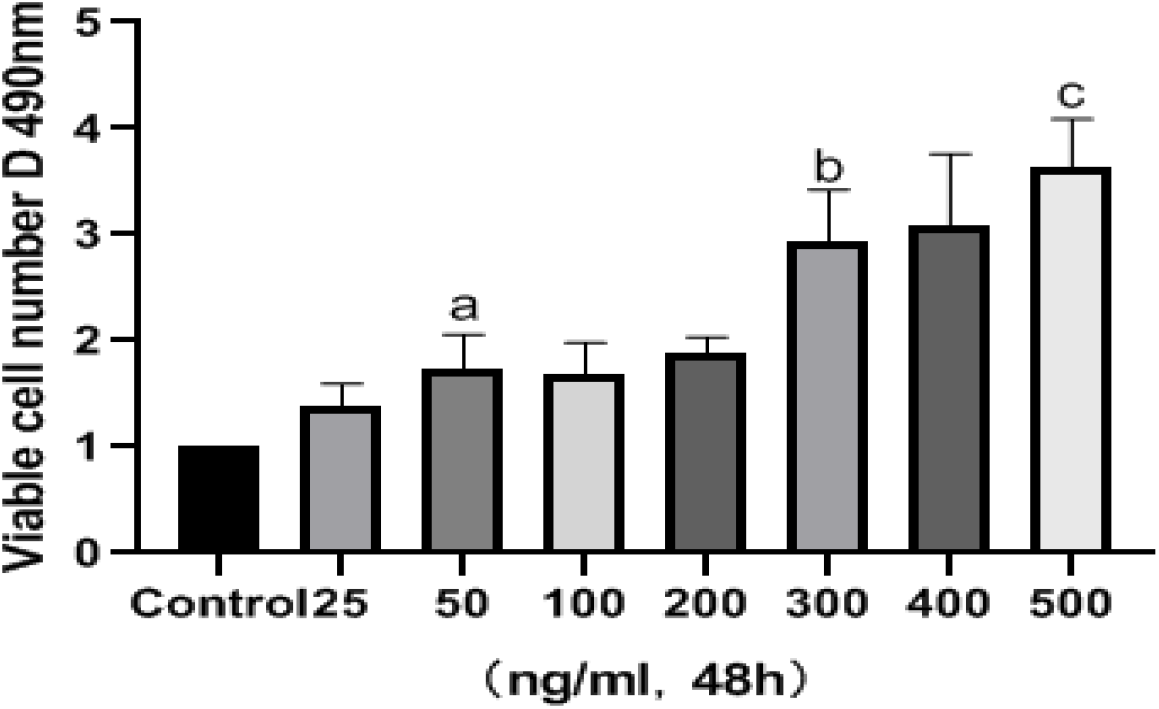
Consequences of varying concentrations of IL-17A on MRC5 cell viability by MTT. Values are mean±SD. *n*=4. *aP*<0.05 *vs* Control group, *bP*<0.05 *vs* 50ng/ml group, *cP*<0.05 vs 300ng/ml group.

### 3. Time dependent effect of 300 ng/ml IL-17A on MRC5 cell proliferation

MRC5 cells were exposed to 300 ng/ml IL-17A for 24 h, 48 h and 72 h, respectively. MTT assay revealed that IL-17A-treated cells had remarkably higher proliferation rate than non-IL-17A treated cells (*aP, bP* and *cP*< 0.05) at 24 h, 48 h and 72 h of treatment. Hence, IL-17A-stimulated MRC proliferation increased significantly, with the prolongation of IL-17A treatment (*eP, fP*< 0.05) (Figure 3).

**Fig. 3.**
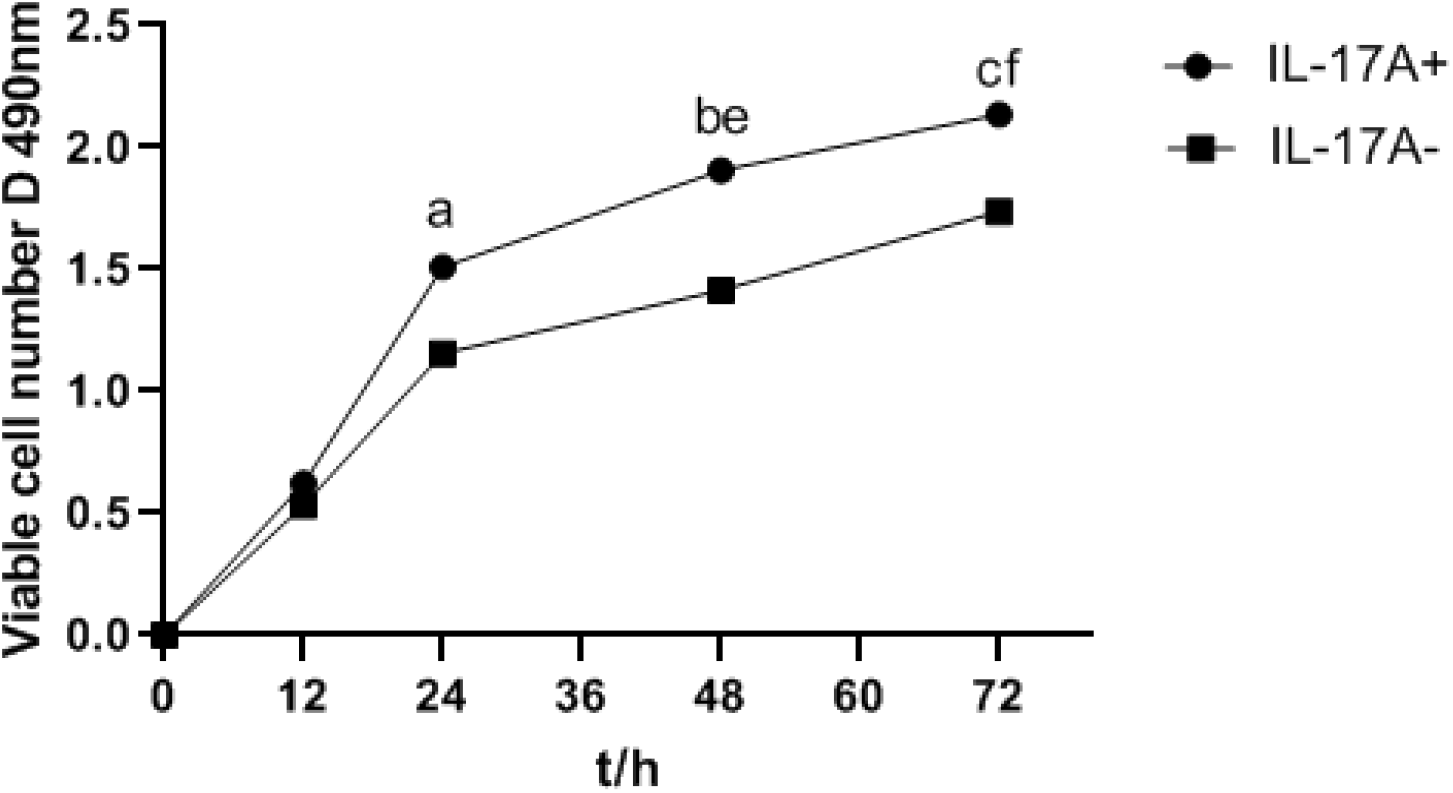
Time dependent effect of 300 ng/ml IL-17A on MRC5 cell proliferation

### 4. Effects of differing concentrations of TSA on MRC5 cell proliferation

Using MTT assay, we revealed that the application of 200 nmol/L TSA significantly decreased MRC5 cell proliferation, indicating that TSA at this concentration had a significant inhibitory effect on the proliferation activity of MRC5 cells (*aP*< 0.05). Moreover, the TSA-mediated suppression of MRC5 cell proliferation became more pronounced with increasing concentrations of TSA (*bP*< 0.05). However, at concentrations of >300 nmol/L, the proliferation suppression was lost (*cP*< 0.05) (Figure 4).

**Fig. 4.**
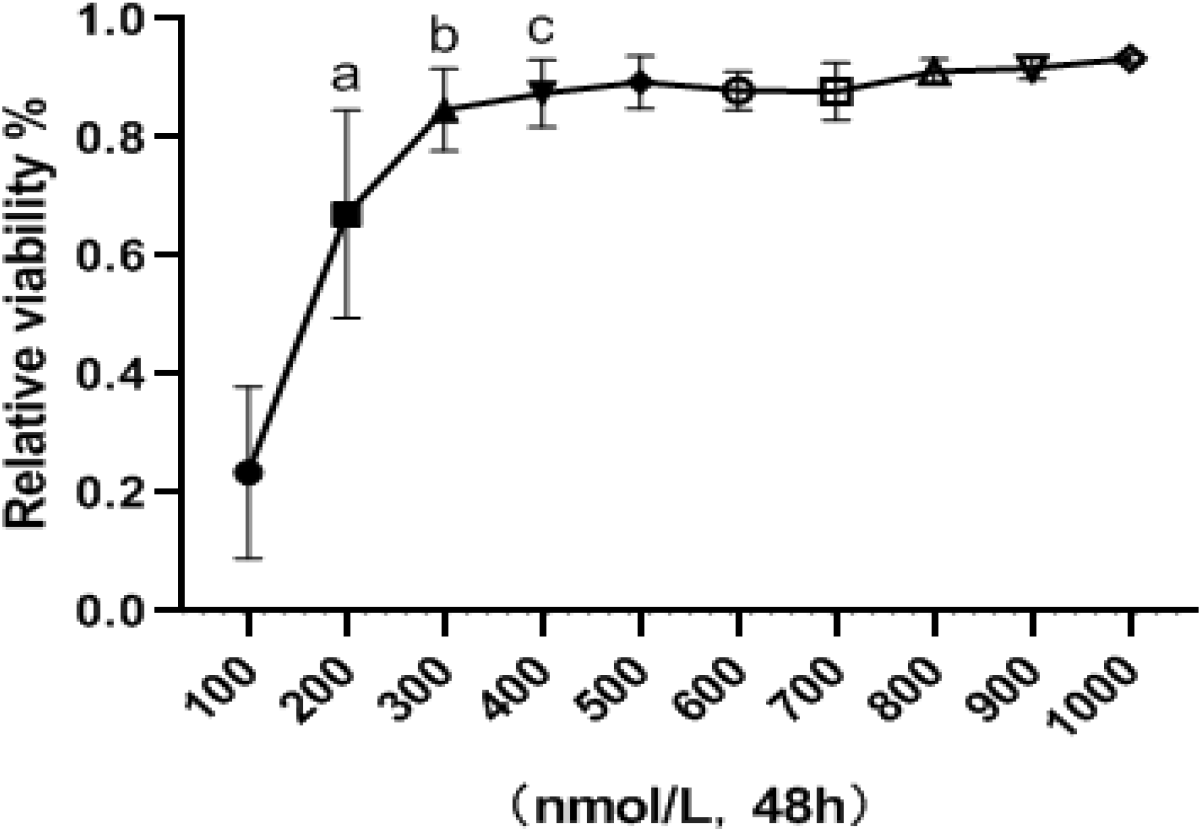
Effects of differing concentrations of TSA on MRC5 cell proliferation

### 5. Time dependent effect of 200 nmol/L TSA on MRC5 cell proliferation

MTT assay examining the suppressive role of 200 nmol/L TSA on MRC5 cell proliferation, based on time, revealed that TSA exposure increased the suppression of cell proliferation over time (*aP*< 0.05). In particular, the suppressive effect of 500 nmol/L TSA on MRC cell proliferation was significantly greater than that of 200 nmol/L TSA (*bP*< 0.05), and it was most obvious at 48 hours. Moreover, the effect of concentrations >500 nmol/L on the proliferative activity of MRC5 cells was not significantly altered. (Figure 5)

**Fig. 5.**
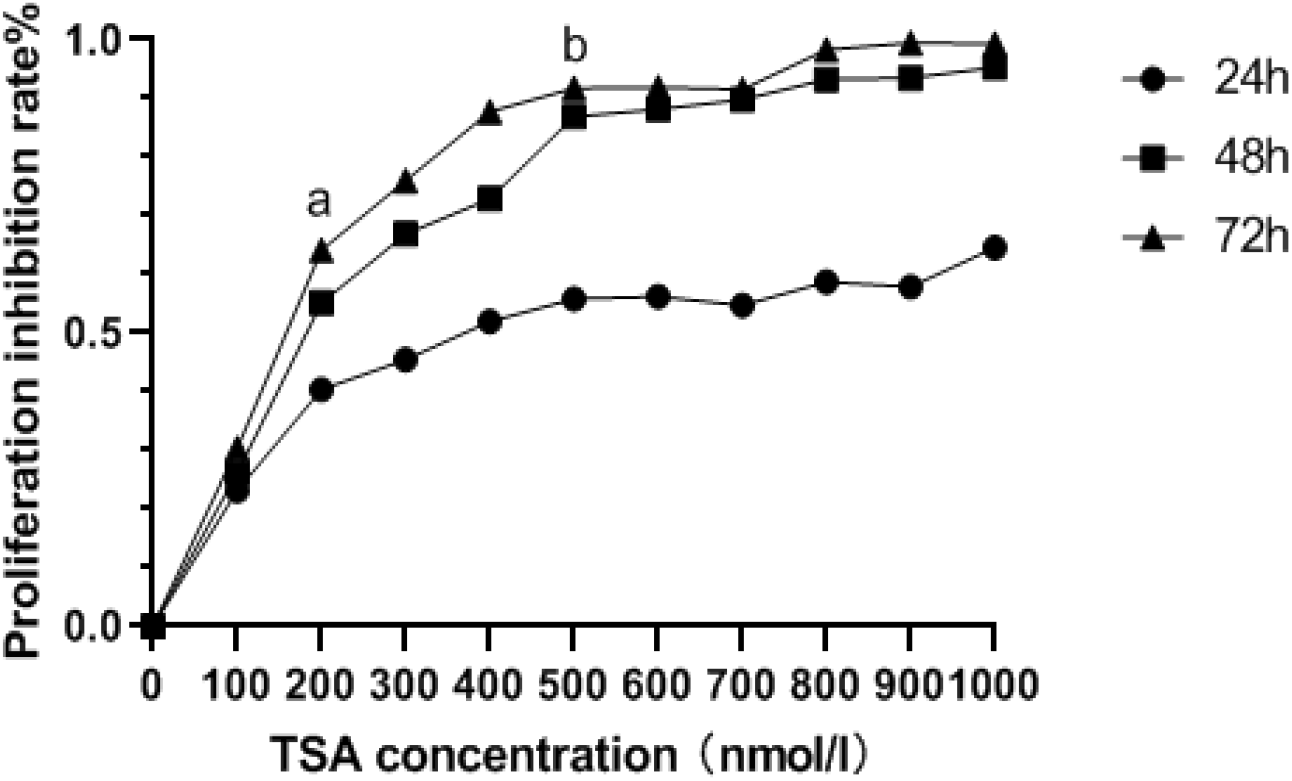
Time dependent analysis of consequences of differing concentrations of TSA

### 6. Results of grouping MRC5 cells

Using microscopic evaluations, we revealed that the IL-17A-treated MRC5 cells experienced markedly elevated rate of proliferation, compared to the controls, and the proliferation density was also increased. Conversely, the growth activity of the TSA-treated MRC5 cells was decreased, and proliferation was significantly inhibited. (Figure 6).

**Fig. 6.**
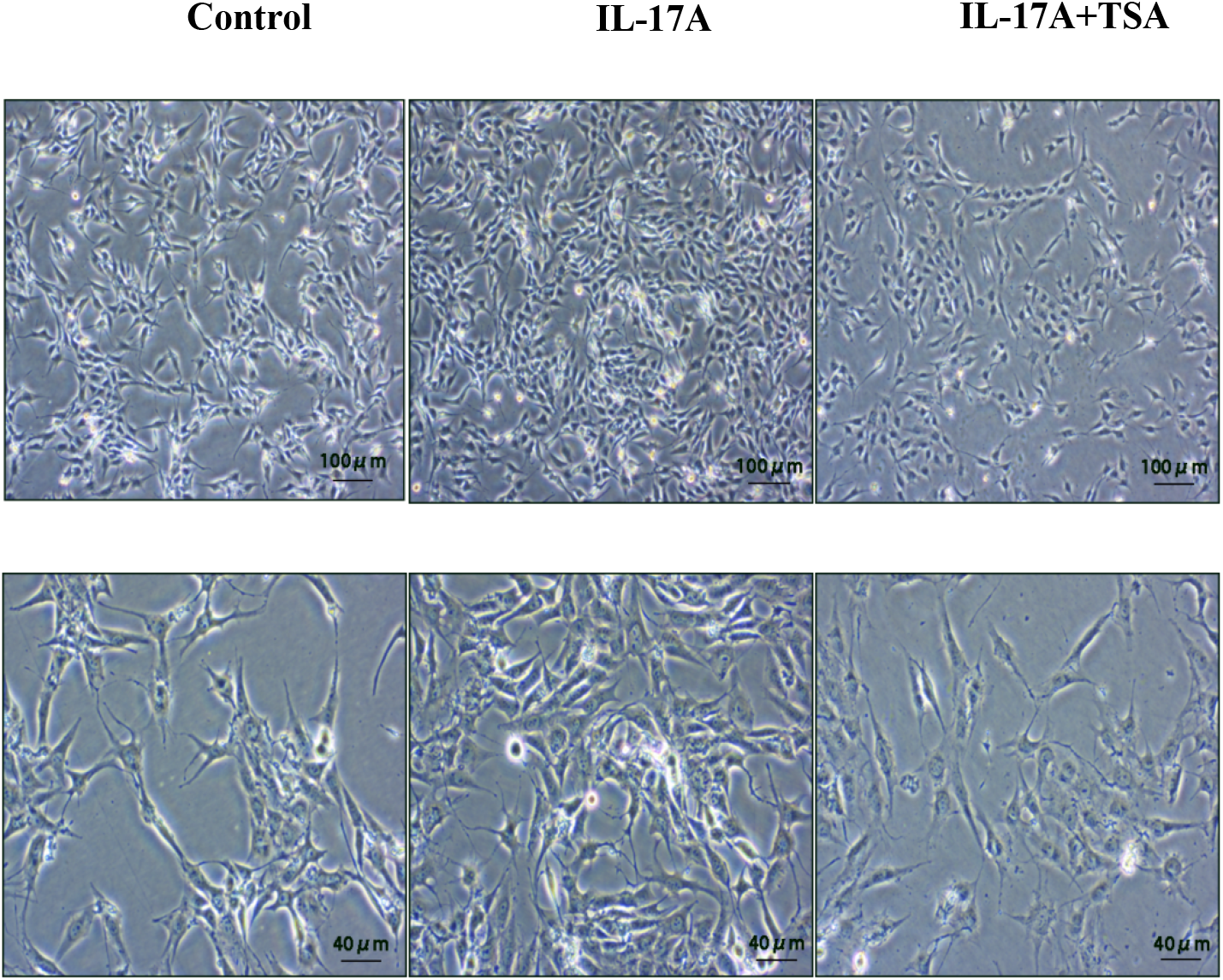
MRC5 cells treated with IL-17 and TSA

### 7. Evaluation of HDAC1 activity

With increasing IL-17A concentration, the HDAC1 activity in MRC5 cells gradually increased. At 50 ng/ml of IL-17A, the HDAC1 activity was markedly elevated, relative to controls (a*P* <0.05). At concentration of 300 ng/ml, the HDAC1 activity was markedly more than the 50 ng/ml IL-17A-treated cells (b*P*< 0.05). At concentration of 500 ng/ml, the HDAC1 activity was markedly increased, relative to controls (a*P*< 0.05). No discernible difference was observed between IL-17A group and 300 ng/ml group (c*P*> 0.05) (Figure 7).

**Fig. 7.**
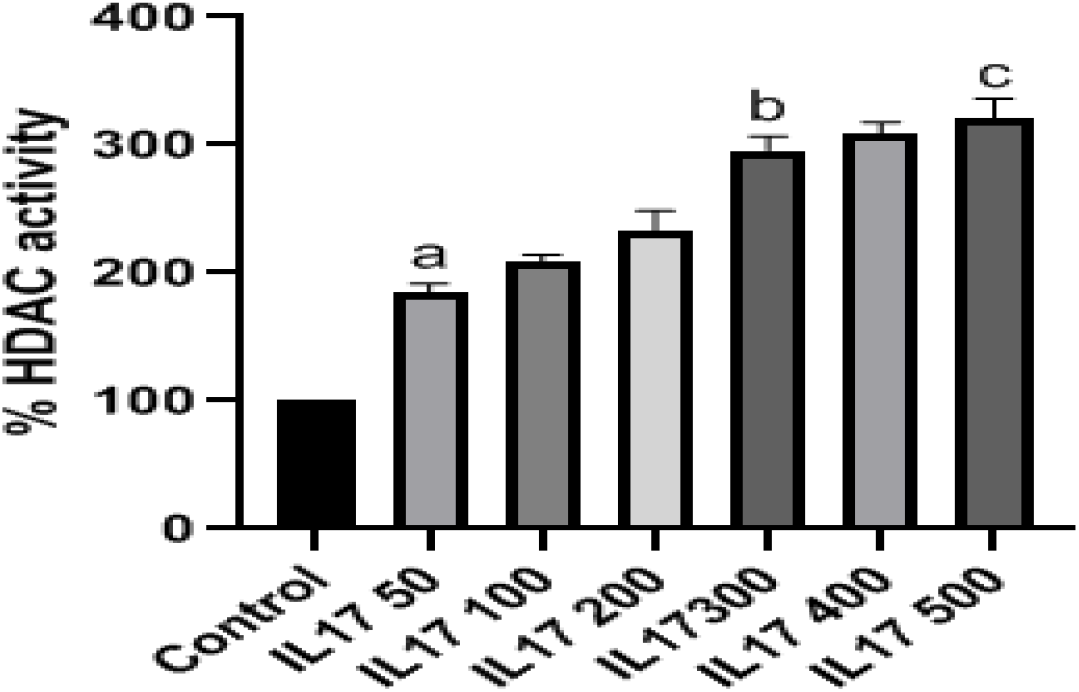
Evaluation of MRC5 cell HDAC1 activity after exposure to varying concentrations of IL-17A

### 8. MRC5 cell HDAC1 activity under differing treatment conditions

MRC5 cells were assigned to five groups, namely control, 50ng/ml IL-17A, 50ng/ml IL-17A + 100nmol/L TSA, 300ng/ml IL-17A and 300ng/ml IL-17A and 200nmol/L TSA. Based on our observations, MRC5 cell HDAC1 activity markedly increased with IL-17A treatment alone verses control (aP, cP < 0.05), and markedly decreased with TSA treatment alone (bP < 0.05 vs IL-17A 200nmol / L, dP < 0.05 vs IL-17A 300 ng / ml group) (Figure 8).

**Fig. 8.**
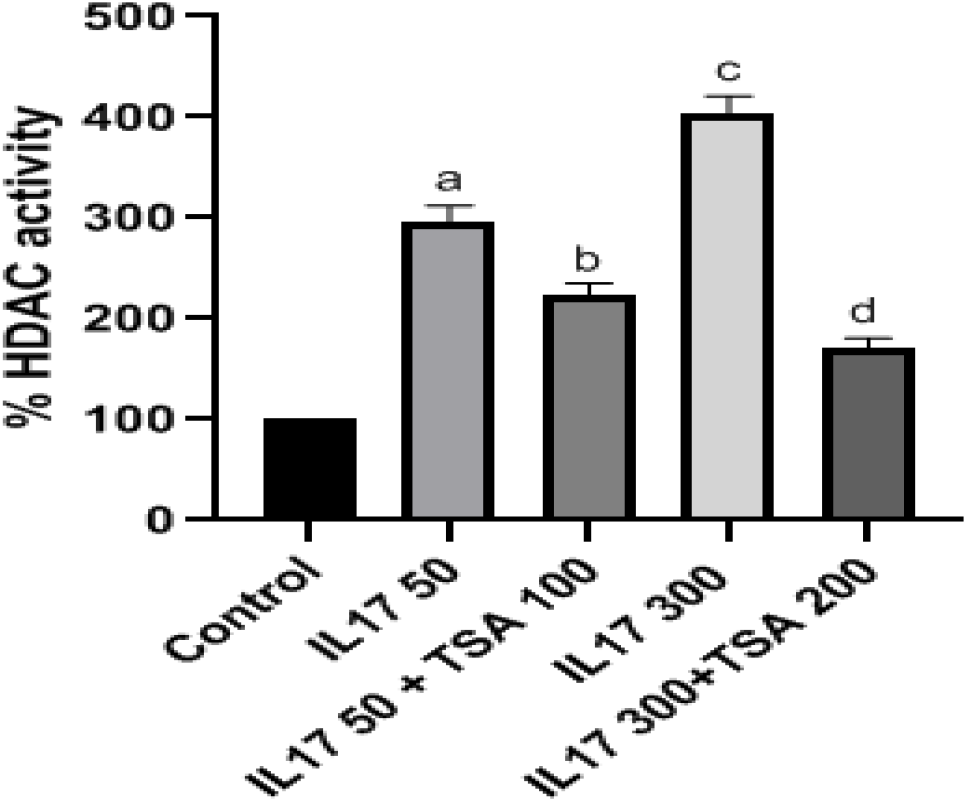
Alterations in HDAC1 activity in MRC5 cells simultaneously exposed to IL-17A and TSA Values are mean±SD. n=4. HDAC1 activity at 50ng/ml was 296.09±16.09 U/mg, *aP*<0.05 VS Control, *bP*<0.05 VS IL-17 50ng/ml group, *cP*<0.05 VS IL-17 50ng/ml group, *dP*>0.05 VS IL-17 300ng/ml group.

### 9. IF evaluations

Relative to controls, Vimentin, Collagen-I and a-SMA levels in the IL-17-stimulated MRC5 cells were significantly increased. Conversely, the levels of the same proteins in the IL-17- and TSA-stimulated MRC5 cells were significantly decreased, compared to the IL-17-stimulated cells, but still remained higher than controls. Of note, Vimentin, Collagen-I and a-SMA levels in control cells were either non-existent or seldomly expressed (Figure 9).

**Fig. 9.**
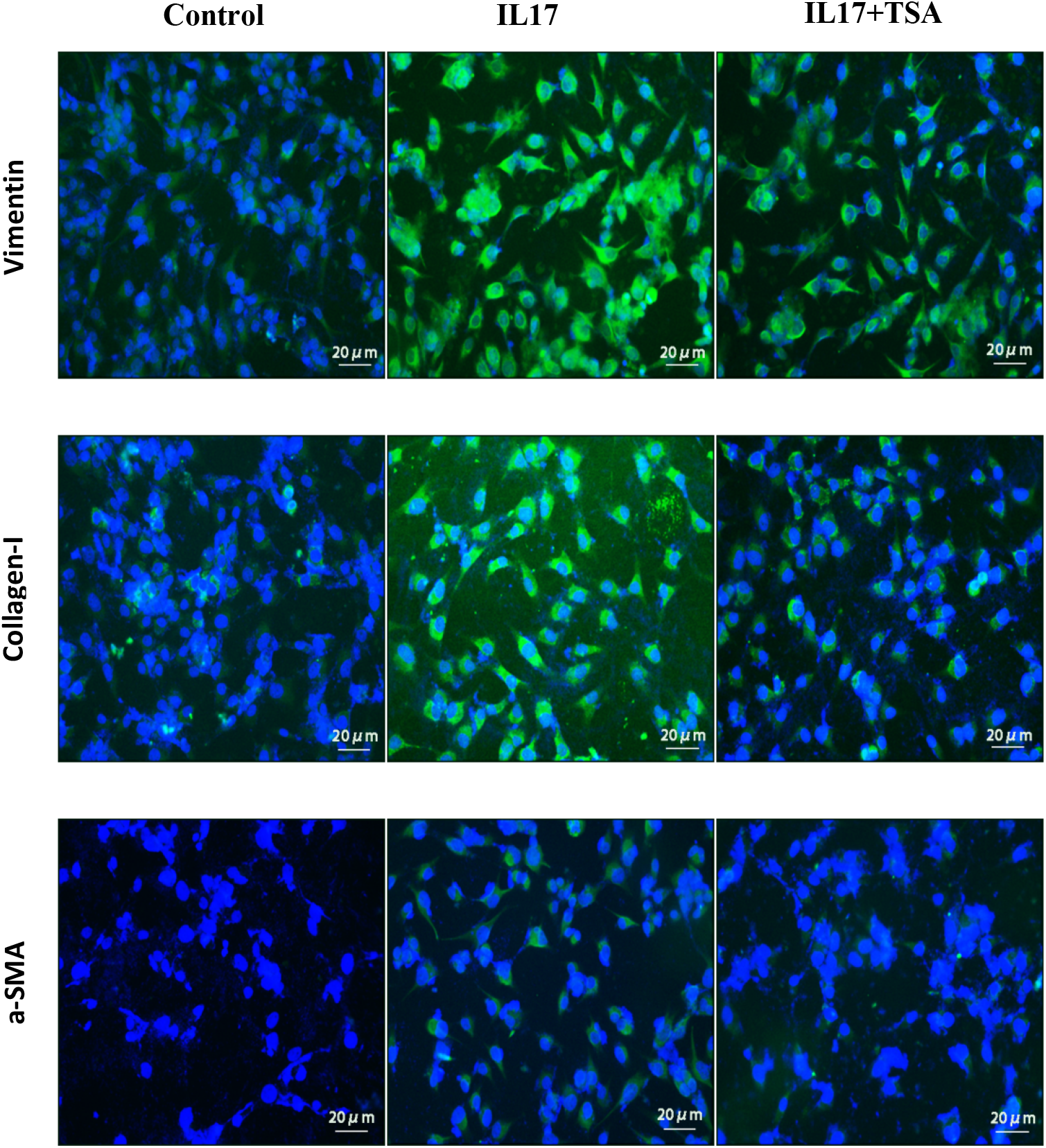
Immunofluorescence staining of Vimentin, Collagen-I and a-SMA in MRC5 cells under differing treatment conditions (×200)

### 10. WB analysis

A-SMA and Vimentin protein levels in the IL-17A-stimulated MRC5 cells were markedly elevated, compared to control cells (*αP*< 0.001). Alternately, the same proteins in the IL-17A- and TSA-stimulated cells experienced a remarkable reduction, compared to IL-17A-stimulation alone (*bP*< 0.001). (Figure 10)

**Fig. 10.**
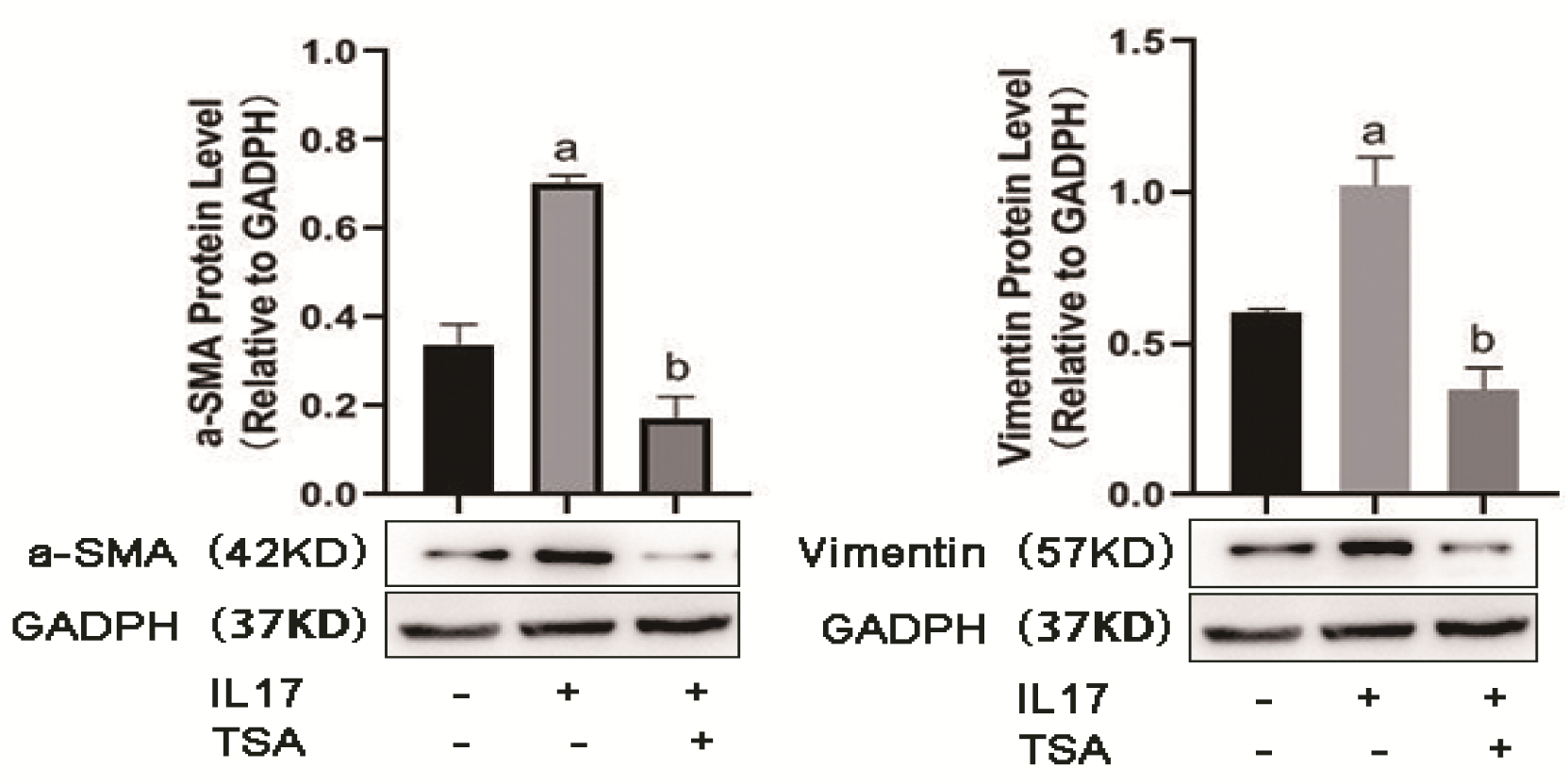
Evaluation of α-SMA and Vimentin protein levels by western blot in MRC5 cells under differing treatment conditions. Values are mean±SD. *n*=4. *aP*< 0.001 *vs* contorl group, *bP*< 0.001*vs IL 17* group.

HDAC1 levels in IL-17A-stimulated MRC5 cells were remarkably higher than in control cells (*aP*< 0.001). Alternately, the levels of the same protein were markedly diminished in cells treated with IL-17 and TSA, relative to IL-17A-stimulated MRC5 cells (*bP* < 0.001). Smad2 levels were lower in the IL-17A- and TSA-stimulated MRC5 cells, relative to controls, however, significance was not achieved (*cP*> 0.05). No discernible difference was observed in the Smad2 levels in MRC5 cells treated with IL-17A and TSA (*dP*> 0.05) (Figure 11).

**Fig. 11.**
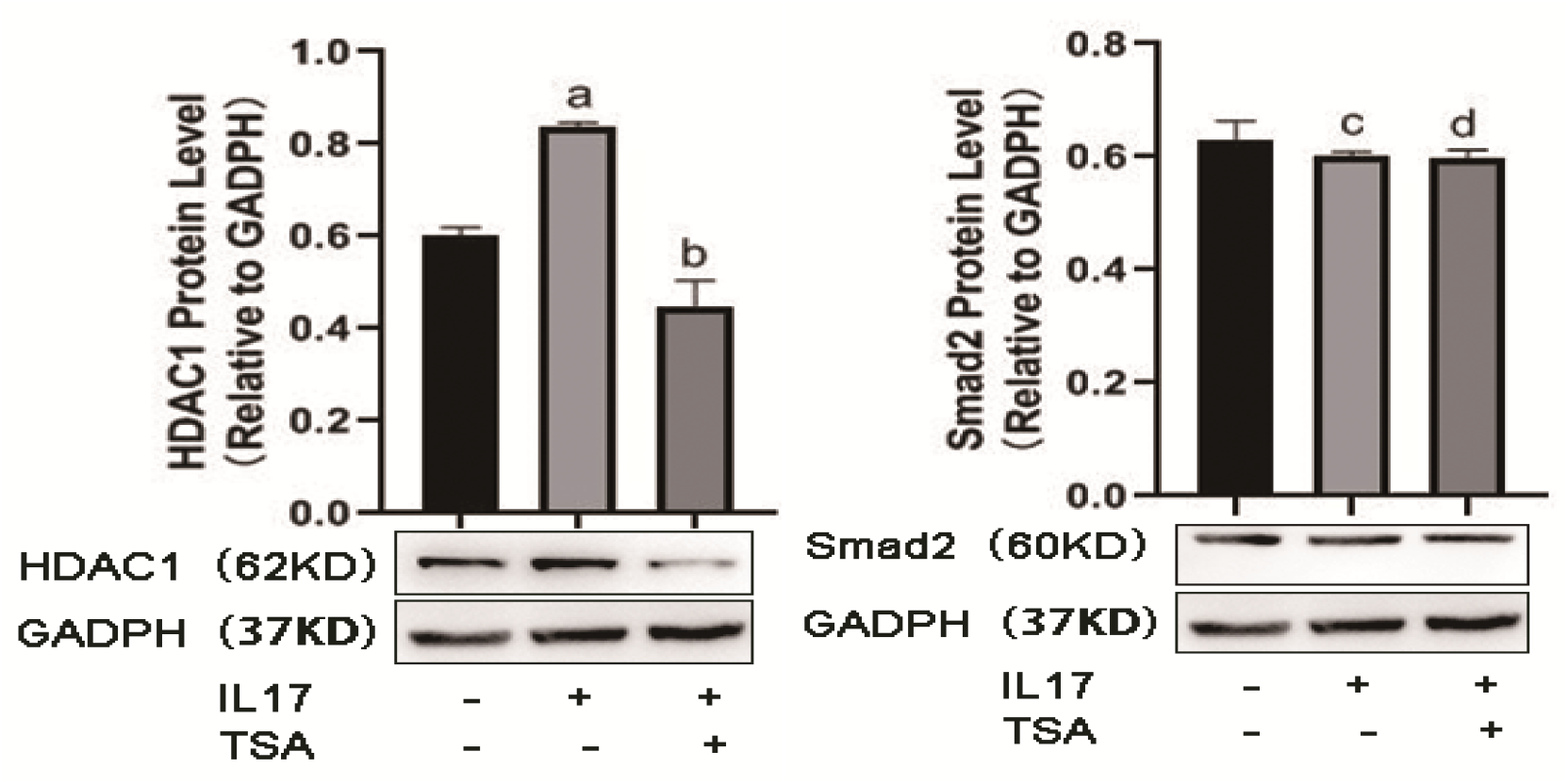
Evaluation of HDAC1 and Smad2 protein levels by western blot in MRC5 cells under differing treatment conditions. Values are mean±SD. *n*=4. *aP*< 0.001 *vs* contorl group, *bP*< 0.001*vs IL 17* group, *cP*>0.05 VS contorl group, *dP*>0.05 VS *IL 17* group.

Smad3 levels in MRC5 cells stimulated by IL-17A alone did not show any significant change, relative to control cells (*cP*> 0.05). Smad3 expression diminished slightly in cells treated with IL-17 and TSA, relative to IL-17 alone, however, it was not significant (*dP*> 0.05). Conversely, Smad7 levels in MRC5 cells stimulated by IL-17A alone was markedly lower, relative to controls (*αP*< 0.001). Furthermore, Smad7 levels in MRC5 cells treated with IL-17 and TSA was remarkably high, compared to IL-17 treatment alone (*bP*< 0.05) (Figure 12).

**Fig. 12.**
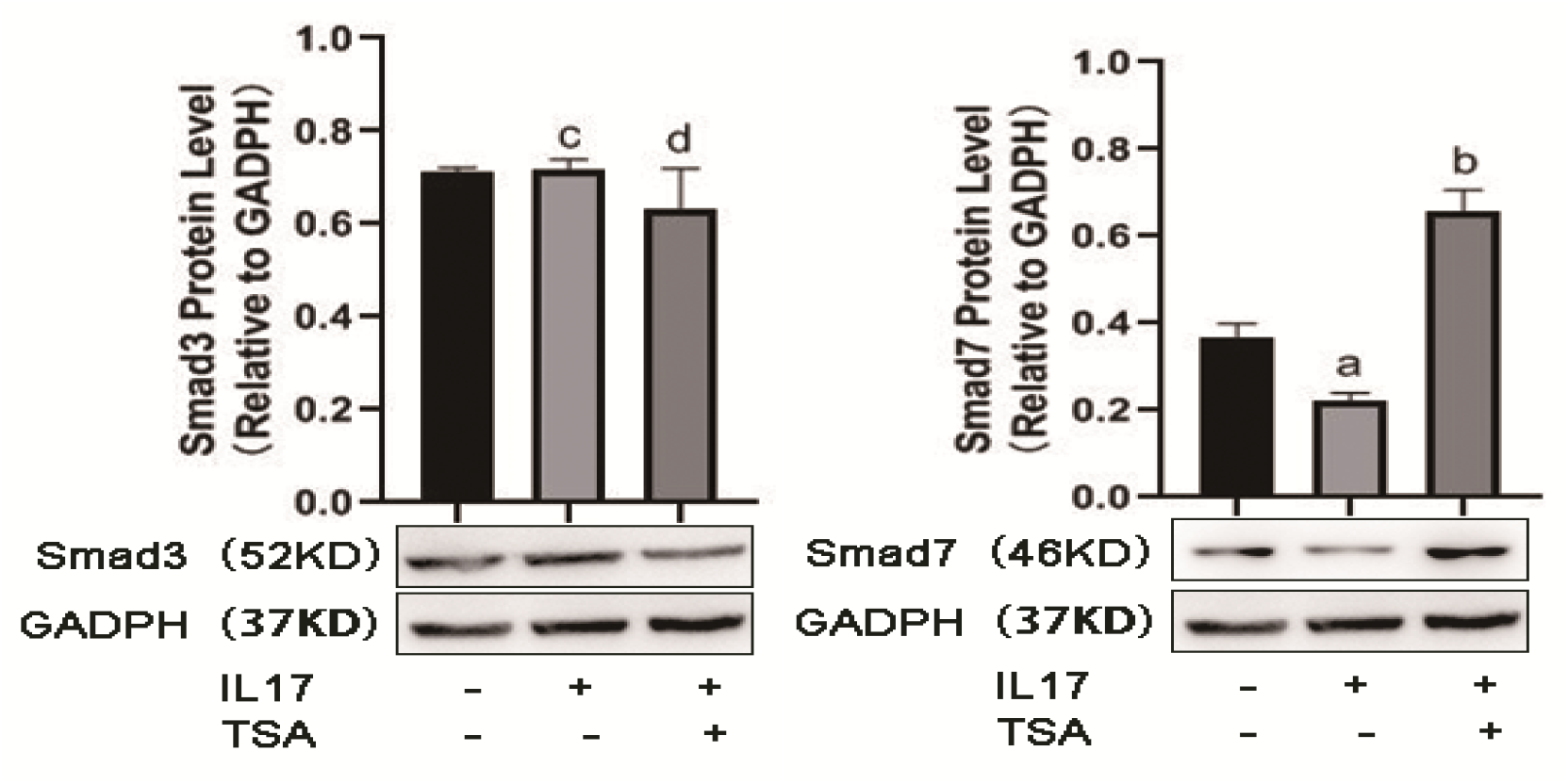
Evaluation of Smad3 and Smad7 protein levels by western blot in MRC5 cells under differing treatment conditions. Values are mean±SD. *n*=4.*cP*>0.05 VS contorl group, *dP*>0.05 VS *IL 17* group, *αP*< 0.001 *vs* contorl group, *bP*< 0.05*vs IL 17* group.

P-Smad2 and p-smad3 protein levels in MRC5 cells stimulated by IL-17A alone was remarkably higher, compared to controls (*aP*< 0.05). Simultaneously, in cells co-treated with IL-17 and TSA, p-Smad2 and p-smad3 levels diminished dramatically, relative to IL-17-treatment alone (*bP*< 0.001) (Figure 13).

**Fig. 13.**
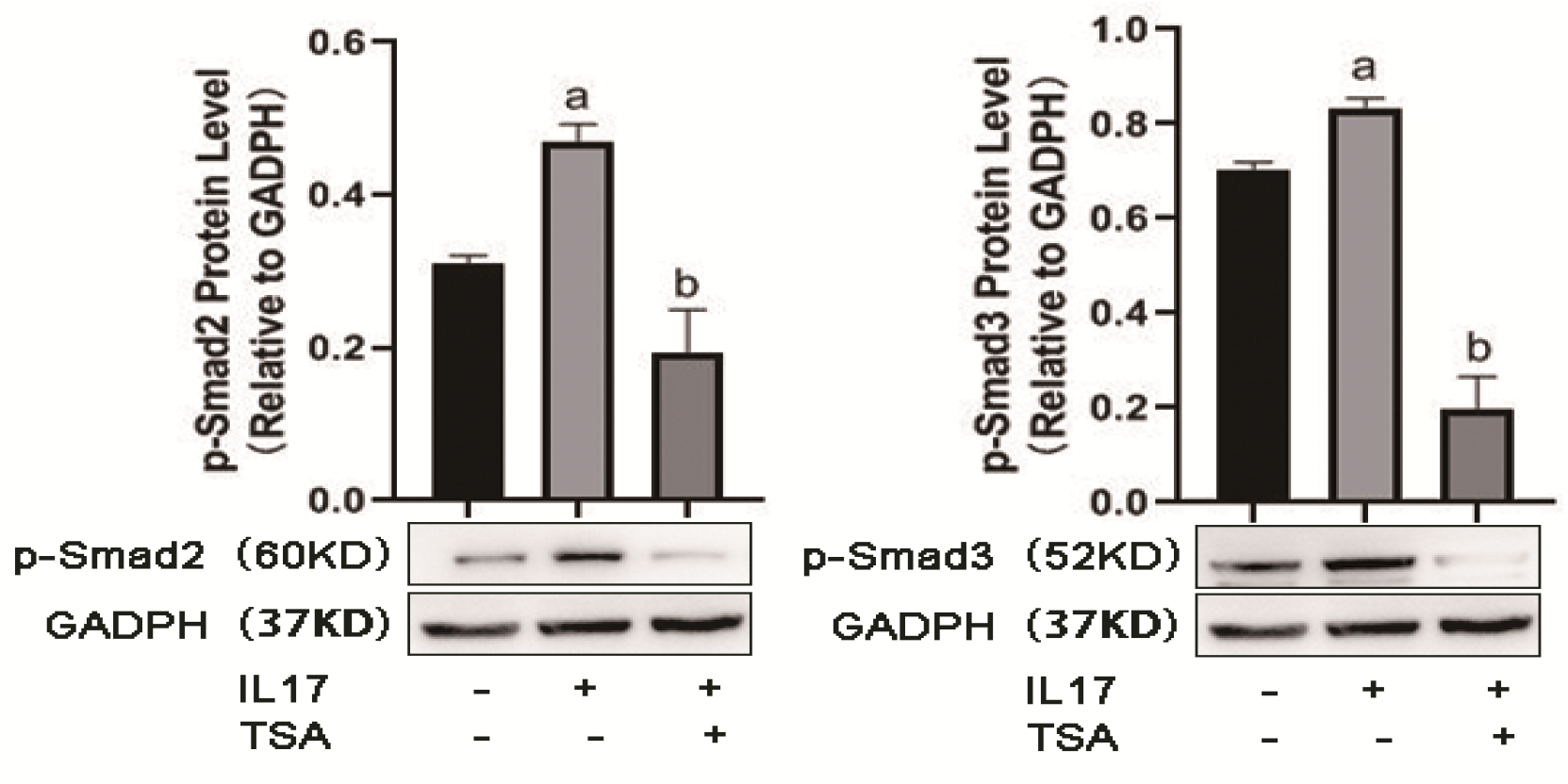
Evaluation of p-Smad2 and p-Smad3 protein levels by western blot in MRC5 cells under differing treatment conditions. Values are mean±SD. *n*=4. *aP*< 0.05 *vs* contorl group, *bP*< 0.001*vs IL 17* group.

## Discussion

Pulmonary fibrosis is a complex process that involves inflammatory cell invasion into lung tissue, conversion of fibroblasts into interstitial cells and the production of a large quantity of interstitial protein deposition that destroys the lung structure. The production and interaction of many cytokines play a key role in this process. In recent reports, it was suggested that IL-17A participates in the occurrence and development of pulmonary fibrosis[1]. In prior experiments, we demonstrated markedly increased Th17 cell differentiation in mice with pulmonary fibrosis. Moreover, the IL-17A levels were markedly upregulated in these mice verses controls. This is indicative of IL-17A playing a principle role in lung fiber formation and development. However, not much is known about the mechanism of IL-17A-stimulated fibroblasts transformation. This study was designed to fill this void.

Interleukin-17A (IL-17A) is an inflammatory cytokine produced by Th17 cell subsets, which can not only activate T lymphocytes, but also induce endothelial cells, epithelial cells and fibroblasts to secrete adhesion molecules, granulocyte macrophage stimulating factor, IL-8, IL-6, etc., thus leading to the development of acute inflammation[2]. It is the most crucial Th17 cell-secreting cytokine and was the first of its kind to be discovered. Subsequent discoveries of cytokines belonging to the same family are IL17B-F and so on. Among the various cytokines, IL-17A is essential to the early inflammatory response and participates in the development of numerous acute and chronic inflammatory diseases[3]. It stimulates the release of pro-inflammatory factor TNF-a, chemokine MCP-1 and metalloproteinase (MMP), and cause inflammatory cell infiltration and tissue destruction[4]. Multiple studies have reported that IL-17 can increase the TNF-a-induced IL-6 levels in the airway smooth muscle cells[5], and stimulate fibroblasts and airway epithelial cells to release granulocyte colony-stimulating factor and granulocyte macrophage colony-stimulating factor[6]. Moreover, in rheumatoid arthritis, IL-17 was shown to activate synovial fluid fibroblasts to secrete inflammatory factors, and stimulate colon myofibroblasts to produce inflammatory mediators[7, 8]. Emerging evidences suggest that IL-17A is associated with the development of chronic inflammatory diseases in the lung, such as chronic obstructive pulmonary disease (COPD), cystic fibrosis, bronchial asthma and so on[9]. Hence, it is clear that IL-17A is intimately involved in recruiting inflammatory cells and in promoting synthesis and release of a variety of inflammatory cytokines. Simultaneously, it can cooperate with a variety of inflammatory factors to amplify the inflammatory effect, and participate in the etiology of autoimmune diseases, acute and chronic inflammation, tumor and so on.

HDACi is known to regulate various immune activities in the body. Its positive or negative effects primarily depend on the type and functional state of immune cells. Several studies demonstrated that HDACi can suppress CD4^+^ T cell activity. In the Moreira[10] study, mice were administered CD4^+^ T cells, prior to TSA treatment. They discovered that TSA inhibited the translocation of NF-κB to the nucleus, resulting in reduced CD4^+^ T cell activity over time. Mechanistically, they proposed that TSA suppressed the expression of specific antigens on the surface of CD4^+^ T cells by elevating p21, an inhibitor of cyclin dependent kinase[11]. Based on these results, TSA can cease CD4^+^ T cell proliferation and, thereby, induce anti-tumor immune response of T cells. Furthermore, Donas et al. [12] discovered that TSA can enhance FOXP3 levels in CD4^+^ T cells, thereby, driving proliferation and differentiation of Treg cell subsets and augmenting immunosuppression. Given these evidences, TSA has great potential in curbing autoimmune diseases.

TSA is a highly prevalent HDACi that can effectively suppress both class I and class II HDACs. Through the inhibition of HDACs, TSA can also suppress TGF - β 1-induced Collagen-I levels in myofibroblasts via negative regulation of transforming factor SPL. In mice, abnormal expression of HDAC2 related nucleoprotein can lead to cardiac interstitial fibrosis, the effects of which can be abrogated by HDACi. Huang et al. [13] revealed that the FAS gene expression in IPF fibroblast tissues was downregulated, due to the deacetylation of the Fas gene promoter. Furthermore, TSA could upregulated Fas gene expression and promoted apoptosis induced by the Fas pathway. Similarly, Sanders et al. [14] demonstrated that TSA can upregulate pulmonary myofibroblasts thymocyte differentiation antigen 1 (Thy-1) transcript levels and suppress fibroblasts proliferation in order to promote its anti-fibrotic role. Moreover, studies[15] have reported that cyclooxygenase (COX) expression is downregulated in patients with pulmonary fibrosis, mainly due to the high deacetylation of the COX gene promoter, while HDACi vorinostat and pabistat can reduce the state of deacetylation of the Cox gene promoter, upregulate expression of COX2, and promote the production of the anti fibrotic factor prostaglandin E2 (PGE2).

One of the obstacles of lung fibroblast detection is that it lacks specific antigen expression. Hence, fluorescence staining exclusion method is primarily used for the identification of lung fibroblasts[16, 17]. The first step, in this process, is establishing that the cells are derived from mesenchymal cells, which is evidenced by the presence of Vimentin. The second step excludes presence of other α-SMA-expressing smooth muscle and endothelial cells. The presence of α-SMA-expressing myofibroblasts is indicative to a full transformation of fibroblasts into myofibroblasts. In this study, we first confirmed the fibroblastic nature of our cells. Subsequently, we revealed, using IF and WB that untreated MRC5 fibroblast cells expressed scarce amounts of Vimentin. However, the levels of Vimentin, Collagen-I, and a-SMA drastically increase in the IL-17A-stimulated MRC5 cells verses controls. Conversely, the Vimentin, Collagen-I, a-SMA levels were strongly suppressed in cells co-treated with IL-17A and TSA, relative to IL-17 treatment alone, however, it still remained higher than controls. These results are indicative of MRC5 cells transforming into myofibroblasts, under IL-17 stimulation, and inhibition of this transformation and secretion of interstitial protein by TSA.

Based on our results, IL-17A stimulates MRC5 cell proliferation, in a concentration-dependent manner. However, the increased proliferative capacity slowed down as IL-17A reached 300ng/ml, and the proliferation capacity decreased at concentration of 500ng/ml. To further clarify the relationship between MRC5 cell proliferation and HDAC1 activity, we analyzed HDAC1 activity under differing treatment conditions. We demonstrated that MRC5
 cell HDAC1 activity increased after IL-17A stimulation alone, in a concentration-dependent manner. TSA, on the other hand, suppressed MRC5 cell proliferation in a concentration- and time-dependent manner. TSA showed obvious inhibition of MRC5 cell proliferation at 200 nmol /L, however, no discernible effect was observed at concentration of 500 nmol/L. To further elucidate the IL-17- and TSA-mediated regulation of MRC5 cell HDAC1 activity, we assigned MRC5 cells to five groups: control, 50hg/ml IL-17, 50ng/ml IL-17 + 100nmol/L TSA, 300ng/ml IL-17 and 300ng/ml IL-17 and 200nmol/L TSA. We demonstrated that HDAC1 activity was markedly reduced in the 50ng/ml IL-17-treated cells, relative to the 300ng/ml IL-17-treated cells and the 300 ng/ml IL-17- and 200 nmol/L TSA-cotreated cells. Moreover, the HDAC1 activity was drastically lowered in the 200 nmol/L TSA-treated cells, relative to the 50 ng/ ml IL-17A- and the 100nmol/l TSA-co-treated cells. The HDAC1 activity in cells treated with IL-17A alone increased significantly. This effect was most obvious at 48h after IL-17A exposure, and the effect was least effective after 72h of IL-17A exposure. Hence, we selected 48 h TSA exposure for our subsequent experiments. Upon MRC5 cell stimulation with IL-17A and TSA, the HDAC1 activity reduced drastically, compared to IL-17A treatment alone, but it remained higher than controls. Furthermore, our evaluation of the WB results revealed that the IL-17A-treated cells had markedly higher levels of the HDAC1 protein, compared to controls. Conversely, in IL-17A- and TSA-co-treated cells, the HDAC1 protein levels diminished significantly. Given these results, TSA can reduce HDAC1 activity via regulation of the HDAC1 protein.

Our protein evaluations also revealed that the levels of HDAC1, phosphorylated Smad2 and phosphorylated Smad3 in the IL-17A-stimulated MRC5 cells were markedly elevated than in control cells, whereas Smad7 protein was markedly reduced than in control cells. In cells co-treated with IL-17A and TSA, the levels of HDAC1, phosphorylated Smad2 and phosphorylated Smad3 were drastically less than in IL-17A-treated cells. Moreover, the levels of Smad7 protein were markedly more than in IL-17A-treated cells. On the contrary, Smad2 and Smad3 levels remained the same in all treatment groups. In summary, the above data revealed that in IL-17A-stimulated MRC5 cells, HDAC1 protein levels increased, HDAC1 activity increased, phosphorylated Smad2 and phosphorylated Smad3 increased, and Smad7 expression decreased, indicating that the TGF-β1/smads pathway was activated. Conversely, the TSA-treated MRC5 cells had dramatically less HDAC1 protein levels, decreased HDAC1 activity, decreased levels of phosphorylated Smad2 and phosphorylated Smad3, and elevated Smad7 levels, which is indicative of TSA-mediated inhibition of the TGF-β1/Smads pathway. Furthermore, using co-treatments of IL-17A and TSA, we further demonstrated that TSA can effectively inhibit the IL-17A-mediated stimulation of the TGF-β1/Smads pathway and the eventual transformation of MRC5 into myofibroblast cells. It was previously suggested that TSA inhibits the expression of extracellular matrix components at both the gene and protein levels during its suppression of MRC5 transformation into myofibroblast cells and fibrosis prevention. Several studies have reported [18–20] Smad7 localization in the nucleus and cytoplasm, using immunofluorescence, however, HDAC1 was always reported to be in the nucleus. Based on these evidences and consistent with our results, we propose that HDAC1 may directly interact with Smad7 in the nucleus. Hence, via inhibition of Smad7, the expression of the phosphorylated Smad2 and phosphorylated Smad3 may be promoted, and the subsequent TGF-β1/smads pathway may be activated.

## Conclusion

In summary, our work revealed that IL-17A promotes the proliferation and transformation of human lung fibroblasts into myofibroblast cells. IL-17A also promotes the production of interstitial protein. TSA showed obvious inhibition of human lung fibroblast proliferation and blocked the activation of key transformation pathways. The process and mechanism of pulmonary fibrosis are quite complex. Therefore, blocking IL-17A secretion, action pathway or monoclonal antibody may play a significant role in inhibiting pulmonary fibrosis. The above research findings may provide new insight into the development and treatment of pulmonary fibrosis.

## Declarations

### Ethics approval and consent to participate

Not applicable.

### Consent for publication

Not applicable.

### Availability of data and materials

Not applicable.

### Authors’ contributions

Qian Zhang, Zexin Guo, and Jing Zhang are mainly responsible for experimental operation and data processing, Qian Zhang is the major contributor in writing the manuscript. Wei Zhou, Yueqing Wu, Lixia Dong and Jing Feng are mainly responsible for the supervision and administration of the experimental projects. Shuo Li and Jie Cao are mainly responsible for editing manuscripts. All authors read and approved the final manuscript.

## Acknowledgements

Not applicable.

## Abbreviations

IL-17A: Interleukin-17A
MRC5: Human normal embryonic lung fibroblasts
TSA: Trichostatin A
HDAC1: Tistone deacetylase
MTT: Cell proliferation and cytotoxicity test kit
TGF-β1: Transforming growth factor-β1
IPF: Idiopathic pulmonary fibrosis
HDACi: Histone deacetylation inhibitors
a-SMA: a-Smooth muscle activator protein
IF: Immunofluorescence
WB: Western blotting
IL-8: Interleukin-8
IL-6: Interleukin-6
TNF-a: Tumor necrosis factor-a
MCP-1: Monocyte chemoattractant protein-1
MMP: Matrix metalloproteinase
COPD: Chronic obstructive pulmonary disease
COX2: Cyclooxygenase-2
PGE2: Prostaglandin E2

